# Differential modulation of ventral tegmental area circuits by the nociceptin/orphanin FQ system

**DOI:** 10.1101/776484

**Authors:** Joseph R. Driscoll, Tanya L. Wallace, Kasra A. Mansourian, William J. Martin, Elyssa B. Margolis

## Abstract

The neuropeptide nociceptin/orphanin FQ (N/OFQ) can be released by stressors and is associated with disorders of emotion regulation and reward processing. N/OFQ and its receptor, NOP, are enriched in dopaminergic pathways, and intra-ventricular agonist delivery decreases dopamine levels in the dorsal striatum, nucleus accumbens (NAc), and ventral tegmental area (VTA). We used whole cell electrophysiology in acute rat midbrain slices to investigate synaptic actions of N/OFQ. N/OFQ was primarily inhibitory, causing outward currents in both immunocytochemically identified dopaminergic (tyrosine hydroxylase positive (TH(+)) and non-dopaminergic (TH(−)) VTA neurons (effect at 1 μM: 20 ± 4 pA). Surprisingly, this effect was mediated by augmentation of postsynaptic GABA_A_R currents, unlike the substantia nigra pars compacta (SNc), where the N/OFQ induced outward currents were K^+^ channel dependent. A smaller population, 19% of all VTA neurons, responded to low concentrations N/OFQ with inward currents (10 nM: −11 ± 2 pA). Following 100 nM N/OFQ, the response to a second N/OFQ application was markedly diminished in VTA neurons (14 ± 10% of first response), but not in SNc neurons (90 ± 20% of first response). N/OFQ generated outward currents in medial prefrontal cortex (mPFC)-projecting VTA neurons, but inward currents in a subset of posterior anterior cingulate cortex-projecting VTA neurons. While N/OFQ inhibited NAc-projecting VTA cell bodies, it had little effect on electrically or optogenetically evoked terminal dopamine release in the NAc measured *ex vivo* with fast scan cyclic voltammetry. These results extend our understanding of the N/OFQ system in brainstem circuits implicated in many neurobehavioral disorders.

**Significance statement:** The neuropeptide nociceptin/orphanin FQ (N/OFQ) and its receptor (NOP) are engaged under conditions of stress and are associated with reward processing disorders. Both peptide and receptor are highly enriched in ventral tegmental area (VTA) pathways underlying motivation and reward. Using whole cell electrophysiology in rat midbrain slices we found: 1) NOPs are functional on both dopaminergic and non-dopaminergic VTA neurons; 2) N/OFQ differentially regulates VTA neurons based on neuroanatomical projection target; and 3) repeated application of N/OFQ produces evidence of receptor desensitization in the VTA but not the SNc. These results reveal candidate mechanisms by which the NOP system regulates motivation and emotion.

## Introduction

Nociceptin/Orphanin FQ (N/OFQ) and its receptor (NOP) make up a neuropeptide signaling system de-orphaned in 1995 (Meunier et al., 1995; Reinscheid et al., 1995) that is engaged under conditions of stress (Ciccocioppo et al., 2000; Devine et al., 2001; Fernandez et al., 2004; Green et al., 2007; Green and Devine, 2009; Leggett et al., 2007, 2006; Nativio et al., 2012; Nicholson et al., 2002). The NOP is a G-protein coupled 7-transmembrane domain receptor that canonically signals through Gi/o proteins, post-synaptically activating G-protein coupled inward-rectifying potassium channels (GIRKs), or pre-synaptically reducing probability of neurotransmitter release via inhibition of N-type calcium channels (Hawes et al., 2000; Knoflach et al., 1996; New and Wong, 2002; Vaughan and Christie, 1996). While amino acid sequence homology has led some to categorize the NOP as an opioid receptor (Bunzow et al., 1994; Meunier et al., 1995; Mollereau et al., 1994; Wang et al., 1994), NOP activation is not blocked by naloxone, a non-selective opioid receptor antagonist that was originally used to classify responses as opioid receptor mediated, blocking activation at mu, delta, and kappa opioid receptors (MOPs, DOPs, and KOPs, respectively) (Gintzler et al., 1997; Mogil and Pasternak, 2001; Reinscheid et al., 1996, 1995). Furthermore, the known endogenous opioid peptides (dynorphins, enkephalins, and endorphins) do not bind to the NOP, and N/OFQ does not bind to the MOP, DOP, or KOP (Ma et al., 1997; Meng et al., 1996; Sim et al., 1996). Because of the extensive amino acid sequence homology and these distinct pharmacological properties, N/OFQ and the NOP are most appropriately subclassified as non-classical members of the opioid family (Cox et al., 2015; Toll et al., 2016).

N/OFQ and the NOP are highly enriched in the ventral tegmental area (VTA), dorsal striatum, nucleus accumbens (NAc), medial prefrontal cortex (mPFC), and central nucleus of the amygdala (Berthele et al., 2003; Neal et al., 1999; Parker et al., 2019). The VTA is the major source of dopamine to limbic forebrain regions and plays a key role in brain networks that coordinate motivation and learned appetitive behaviors (Fields et al., 2007). Activity of VTA dopamine neurons is associated with salience and reward prediction, while destruction of these neurons results in motivational deficits (Fields et al., 2007; Kim et al., 2012; Mohebi et al., 2019; Morales and Margolis, 2017; Tsai et al., 2009; Ungerstedt, 1971; Wise, 2005; Witten et al., 2011). Intracerebroventricular (ICV) injections of N/OFQ produce a decrease in extracellular dopamine in the dorsal striatum and NAc, and some midbrain putative dopamine cell bodies are inhibited by NOP activation (Di Giannuario and Pieretti, 2000; Lutfy et al., 2001; Murphy et al., 1996; Murphy and Maidment, 1999; Vazquez-DeRose et al., 2013; Zheng et al., 2002).

Dysregulation of the N/OFQ system has been associated with disorders of motivated responding (Civelli, 2008), and the N/OFQ system has been investigated as a novel therapeutic target for major depressive disorder and alcohol use disorder (Witkin et al., 2019), however understanding the involvement of the N/OFQ system in these behaviors remains a challenge. In fact, in some cases, activation and blockade of NOPs paradoxically produce the same behavioral outcomes, such as with alcohol consumption (Ciccocioppo et al., 2014, p. 7716; Kuzmin et al., 2007; Rorick-Kehn et al., 2016) and anxiety-related behaviors (Dautzenberg et al., 2001; Fernandez et al., 2004; Gavioli et al., 2002; Green et al., 2007; Jenck et al., 1997; Kamei et al., 2004; Varty et al., 2008; Vitale et al., 2006). Such observations may be explained by off-target effects of N/OFQ, activation of N/OFQ sensitive neural circuits that compete for behavioral control, or receptor desensitization.

Here we investigated the basic physiology of N/OFQ responses in VTA neurons to better characterize how N/OFQ contributes to motivation and reward processing. To confirm that our physiological responses to N/OFQ were due to NOP activation we utilized the selective NOP antagonist BTRX-246040 (Toledo et al., 2014) to block N/OFQ responses. We observed similar N/OFQ effects on both dopamine and non-dopamine VTA neurons. Importantly, we found that responses to N/OFQ differ between VTA and substantia nigra pars compacta (SNc) in mechanism of inhibition and functional desensitization measures. Furthermore, we found that for VTA neurons, N/OFQ responses vary by the projection target. For example, N/OFQ induced small inward currents preferentially in VTA neurons that project to the posterior anterior cingulate cortex (pACC). In addition, although NAc-projecting cell bodies were inhibited by NOP activation, N/OFQ did not inhibit dopamine release at terminals in the NAc. Together these observations indicate that NOP actions vary not only by brain region and neuron subpopulation, but also by structural localization within a neuron.

## Materials and Methods

### Electrophysiology

Most experiments were completed in tissue from male Sprague Dawley rats, p22 – p36, except mechanism experiments which were completed in tissue from adult rats (>200g). Rats were anesthetized with isoflurane, and brains were removed. The brains were submerged in Ringer’s solution containing (in mM): 119 NaCl, 2.5 KCl, 1.3 MgSO_4_, 1.0 NaH_2_PO_4_, 2.5 CaCl_2_, 26.2 NaHCO_3_, and 11 glucose saturated with 95% O_2_–5% CO_2_ and horizontal brain slices (150 μm thick) containing the VTA were prepared using a Vibratome (Leica Instruments, Nussloch, Germany). Slices were and allowed to recover at 35°C for at least 1 hr before recordings were initiated. The same Ringer’s solution was used for cutting, recovery, and recording.

Individual slices were visualized under an Olympus BX50WI microscope (Olympus Life Science Solutions, Waltham, MA) with differential interference contrast optics and near infrared illumination, using an Andor xIon+ camera, and Andor Solis imaging software (Andor Technology Ltd, Belfast, Northern Ireland), or under a Zeiss Axio Examiner.D1 with differential interference contrast optics, near infrared illumination, and Dodt contrast, using a monochrome Axiocam 506 (Zeiss International, Oberkochen, Germany). Whole-cell patch-clamp recordings were made at 33°C using 2.5– 4M pipettes containing (in mM): 123 K-gluconate, 10 HEPES, 0.2 EGTA, 8 NaCl, 2 MgATP, and 0.3 Na_3_GTP, pH 7.2, osmolarity adjusted to 275 mOsm. Biocytin (0.1%) was added to the internal solution for post hoc identification.

Recordings were made using an Axopatch 1-D (Axon Instruments, Union City, CA), filtered at 2 kHz, and collected at 20 kHz using IGOR Pro (Wavemetrics, Lake Oswego, OR) or an IPA amplifier with SutterPatch software (Sutter Instrument, Novato, CA) filtered at 1 kHz and collected at 10 kHz. Liquid junction potentials were not corrected during recordings. Hyperpolarization-activated cation currents (*I*_h_) were recorded by voltage clamping cells and stepping from −60 to −40, −50, −70, −80, −90, −100, −110, and −120 mV. The *I*_h_ magnitude was measured as the difference between the initial response to the voltage step after the capacitive peak and the final current response.

Pharmacology experiments were completed in voltage-clamp mode (V = −60 mV) to measure changes in membrane current. Series resistance was monitored online by measuring the peak of the capacitance transient in response to a −4 mV voltage step applied at the onset of each sweep. Input resistance was measured using the steady state response to the same voltage step. Upon breaking into the cell, at least 10 min was allowed for the cell to stabilize and for the pipette internal solution to dialyze into the cell. Drugs were applied via bath perfusion at a flow rate of 2 mL/min or pressure ejection using a SmartSquirt micro-perfusion system (AutoMate Scientific, Berkeley, CA) coupled to a 250 μm inner diameter tubing outlet positioned nearby the recorded cell (within ~200 μm). N/OFQ (1 nM to 10 *μ*M) was bath applied (5-7 min) or pressure injected (2 min) only after a 5 min stable baseline was achieved. Responses were similar to the two forms of N/OFQ application at the same concentrations. For instance at 100 nM, bath application 10.1 ± 1.5 pA, n = 21; pressure ejection 9.8 ± 2.1 pA, n = 12. Any cell that showed drift or did not maintain a consistent baseline current for the full 5 min period was removed from the analysis. All experiments where repeated N/OFQ applications are reported, such as the desensitization experiments, were completed with bath application. To test that observed N/OFQ-mediated effects were specific to NOP, the selective NOP antagonist BTRX-246040 (10 or 100 nM) was applied for 10 min prior to N/OFQ. As there was no statistical difference in the mean amplitude of response for bath application and pressure injection the results were combined for the analysis.

For iontophoresis experiments, the holding current was set to −50 mV to increase the driving potential for Cl^−^. GABA (100 mM, pH adjusted to 4.9 with 37% HCl) was prepared daily and the GABA-containing pipette was positioned approximately 50 μm away from the recorded neuron. Negative retention current (approximately −35 nA) was applied to the GABA pipette, interrupted by positive ejection current pulses (100 ms) once every 30 s, with the intensity adjusted so that the response amplitude was in the range of 100-300 pA.

Stock solutions of drugs were made in advance, stored at −20°C, and diluted into aCSF immediately before application. N/OFQ was obtained from Tocris (Minneapolis, MN) and diluted to a 100 *μ*M stock solution in ddH_2_O. Stock BTRX-246040 was obtained from BlackThorn Therapeutics and dissolved in DMSO (10 mM).

### Retrograde Tracer Injections

Male Sprague Dawley rats, 21–100 d old, were anesthetized with isoflurane. A glass pipette (30- to 50-μm tip) connected to a Nanoject II/Nanoliter 2000 microinjector (Drummond Scientific Co.) was stereotaxically placed in the mPFC (from bregma [in mm]: anteroposterior [AP], +2.6; mediolateral [ML], ±0.8; ventral [DV], −4.0 from skull surface), the pACC (AP, 1.6; ML, ± 0.6; V, −3.5), or the NAc (AP, +1.5; ML, ± 0.8; V, −6.7). Neuro-DiI (7% in ethanol; Biotium) was slowly injected, 50.6 nL per side. Animals were allowed to recover for 5 to 7 days while the retrograde tracer transported back to the cell bodies. On the day of recording, the experimenter was blind to the location of retrograde tracer injection (mPFC, pACC, or NAc) and slices were prepared as above. Projection neurons were chosen by selecting cells observed as labeled using epiflorescent illumination. All injection sites were histologically confirmed by a third party blind to the electrophysiology results to avoid bias. N/OFQ responses were analyzed prior to unblinding. Animals with improper injection placements or significant diffusion outside of the target region were rejected.

### Immunohistochemistry

Slices were pre-blocked for 2 h at room temperature in PBS with 0.2% BSA and 5% normal goat serum, then incubated at 4°C with a rabbit anti-TH polyclonal antibody (1:100; EMD Millipore, RRID: AB_390204). Slices were then washed thoroughly in PBS with 0.2% BSA before being agitated overnight at 4°C with Cy5 anti-rabbit secondary antibody (1:100; Jackson ImmunoResearch Labs Inc., West Grove, PA, RRID: AB_2534032) and FITC streptavidin (6.5 μL/mL). Sections were rinsed and mounted on slides using Bio-Rad Fluoroguard Antifade Reagent mounting media and visualized with an Axioskop FS2 Plus microscope with an Axiocam MRm running Neurolucida (MBF Biosciences, Williston, VT). Neurons were only considered TH(−) if there was no colocalization of biocytin with TH signal and the biocytin soma was in the same focal plane as other TH(+) cell bodies. Primary antibodies were obtained from Millipore Bioscience Research Reagents or Millipore, secondary antibodies were obtained from Jackson ImmunoResearch Laboratories, and all other reagents were obtained from Sigma Chemical.

### Fast Scan Cyclic Voltammetry

Male Sprague Dawley rats, 21–26 d old, or *Th∷Cre* transgenic rats (Witten et al., 2011), 46-51 d old at the time of virus injection, were used in these studies. Th∷Cre rats were injected with the Cre dependent ChR2 expressing virus (AAV2-Ef1a-DIO-hChR2(H134R)-mCherry, titer 5.1×10^12^ viral particles/mL, UPenn viral core) bilaterally into the VTA 500 nL per side (AP, −5.3; ML, ± 0.4; DV, −8.2 mm from bregma). Five weeks later, coronal slices (400 μm) containing the NAc were prepared for voltammetry measurements. The use of Cre dependent ChR2 expression allowed selective optical control of VTA dopamine terminals in the NAc.

Extracellular dopamine release was achieved using either electrical (in wild-type Sprague Dawley rats) or 470 nm light (in Th∷Cre rats) stimulation. Stimulation parameters were the same for both electrical and optical stimulation (10 Hz, 2 pulses, 4 ms). Electrochemical recordings were made using carbon fiber electrodes fabricated from T-650 carbon fiber (7 μm diameter, gift from Dr. Leslie Sombers (NCSU)) that was aspirated into a borosilicate glass capillary (0.6 × 0.4 mm or 1.0 × 0.5 mm diameter, King Precision Glass Inc., Claremont, CA) and pulled using a PE-22 puller (Narishige, Tokyo, Japan). Carbon fiber electrodes were positioned 80 μm into the tissue either between the bipolar tips of the stimulating electrode or directly in front of an optical fiber connected to an LED emitting 470 nm light (7-10 mW). The potential of the carbon fiber electrode was held at −0.4 V relative to the Ag/AgCl reference electrode. A triangle wave form was passed through the carbon fiber driving the potential from −0.4 V to +1.3 V and back to −0.4 V at a rate of 400 V/s, at 60 Hz for conditioning and 10 Hz for data collection. Data were collected with a WaveNeuro fast scan cyclic voltammetry (FSCV) potentiostat (Pine Research, Durham, NC) using HDCV acquisition software package (freely available through UNC Department of Chemistry). HDCV Acquisition Software was used to output the electrochemical waveform and for signal processing (background subtraction, signal averaging, and digital filtering (4-pole Bessel filter, 2.5 kHz)). Dopamine release was stimulated at 2 min intervals for electrical stimulation and 3 min intervals for optical stimulation. The difference in stimulation invervals was to decrease rundown of the dopamine release signal that can be particularly strong in optical experiments as reported in (Bass et al., 2013; O’Neill et al., 2017). Mean background currents from 1 sec of data prior to stimulation were removed by subtraction of cyclic voltammograms for each trial.

### Data Analysis

For electrophysiology, effects of N/OFQ were statistically evaluated in each neuron by binning data into 30 s data points and comparing the last eight binned pre-drug points to the last eight binned points during drug application using Student’s unpaired *t* test. To evaluate the output of this analysis approach, we performed a subsequent sliding window analysis on this classified data from TH(+) neurons that were tested with 10 nM N/OFQ (Figure 1-1). The results of this analysis are consistent with this classification scheme identifying drug responses and a lack of contamination by drift in individual recordings. For within cell comparisons of N/OFQ responses, responses were compared with a Student’s paired *t* test. *P* < 0.05 was required for significance in all analyses. Differences between neuron populations were tested using two-tailed permutation analyses unless otherwise indicated. Violin plots were constructed by calculating the kernal density estimate, made using a Scott estimator for the kernal bandwidth estimation. The kernel size was determined by multiplying the Scott bandwidth factor by the standard deviation of the data within each bin. Each individual violin plot was normalized to have an equal area under the curve. Time course figures are averages of the binned current traces for all cells time locked to the start of drug application. EC_50_ was estimated by fitting the concentration response data with the Hill equation. Results are presented as mean and standard error of the mean (SEM). Custom code created for analyses here are publicly available at https://osf.io/c8gu7/?view_only=63ea4c0623b54e46a4efaccc450a89c6.

**Figure 1:**
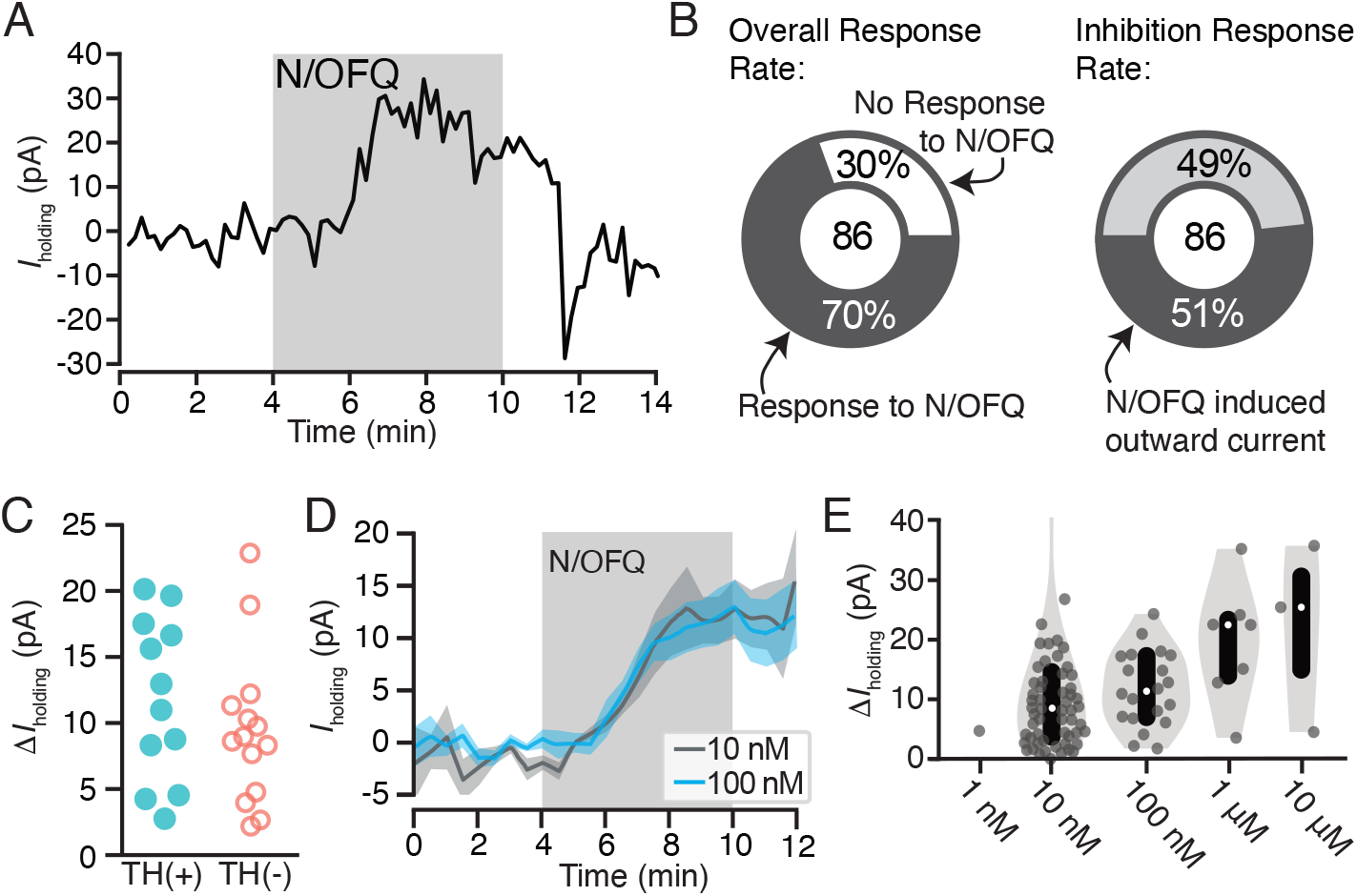
N/OFQ induced outward currents in a subset of VTA neurons. A: Example voltage clamp recording (V_clamp_ = −60 mV) of a VTA neuron that responded to N/OFQ with an outward current. B: Across recordings in neurons from control rats, the majority of VTA neurons responded to 10 nM N/OFQ application (60 out of 86 neurons responded). Forty four out of 60 responses were outward currents. C: A subset of recorded neurons were recovered following whole cell recording and immunocytochemically identified for TH content, a marker for dopamine neurons. Outward currents of similar magnitudes were observed in TH(+) and TH(−) neurons. D: The time courses and maximal effects of bath application of 10 nM and 100 nM N/OFQ were similar. E: Concentration response relationship for VTA neurons showing a positive change, both significant and not significant, in holding current with N/OFQ application (grey dots include all neurons with a change > 0 pA; median shown in white dots; black bars show 25 and 75 percentiles; 1 nM: n = 1/6; 10 nM: n = 55/86; 100 nM: n = 20/25; 1 μM: n = 7/7; 10 μM: n = 3/3).

## Results

### N/OFQ effects on holding current in VTA dopamine and non-dopamine neurons

To test the postsynaptic responses of VTA neurons to N/OFQ, we made *ex vivo* whole cell voltage clamp recordings (V_m_ = −60 mV). N/OFQ application changed the holding current in 70% (60/86) of neurons tested in the VTA (10 nM; 86 neurons from 59 rats; Fig. 1A,B). The majority of responses were relatively small outward currents (73% of responsive neurons, 44/60; 51% of all neurons tested, 44/86; mean response magnitude = 15 ± 2 pA; Fig. 1D). In many cases the holding current returned to baseline during N/OFQ washout, as in Fig. 1A, however in some cases we observed only partial recovery. Using post-hoc immunocytochemistry, we analyzed TH content in each histologically recovered neuron and found that N/OFQ inhibited both confirmed dopamine and non-dopamine neurons in similar proportions (of 44 inhibited neurons from 38 rats, 26 neurons from 23 rats were identified: TH(+): 12/26; TH(−): 14/26). The magnitudes of responses were also similar between confirmed dopamine and non-dopamine neurons (TH(+): 12 ± 2 pA (n = 12); TH(−): 9 ± 2 pA (n = 14); *p* = 0.3 two tailed permutation test; Fig. 1C). The EC_50_ for these outward currents is in the nM range (8 ± 6 nM; Fig. 1E).

To confirm responses were due to activation of the NOP, we tested whether these inhibitions were blocked by the selective NOP antagonist BTRX-246040. In neurons responding to N/OFQ with an outward current (10 nM mean response = 14 ± 3 pA) BTRX-246040 (100 nM) was applied for 10 min and then N/OFQ was applied again in the presence of the antagonist. BTRX-246040 consistently and completely blocked N/OFQ-induced outward currents (baseline N/OFQ response: 14 ± 3 pA; N/OFQ response in BTRX-246040: −1 ± 2 pA; n = 15; 14 rats; paired t-test: *p* = 0.0005; Fig. 2).

**Figure 2:**
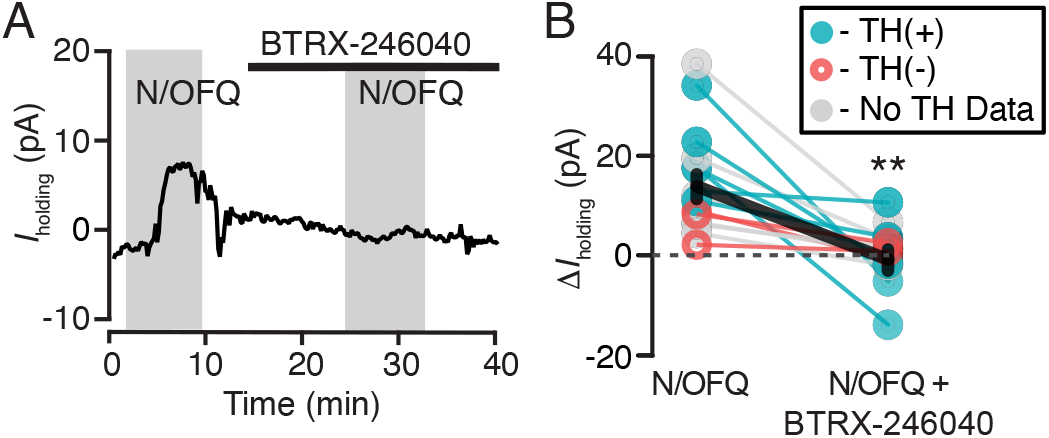
BTRX-246040 consistently blocks N/OFQ induced currents. A: Example recording of a N/OFQ (10 nM) responsive neuron where the selective NOP antagonist BTRX-246040 (100 nM) blocked the response to a subsequent N/OFQ application. B: BTRX-246040 blocked N/OFQ responses across VTA neurons, including both TH(+) and TH(−) neurons (n = 6 and 3, respectively; n = 6 no TH data; mean ± SEM in black). \*\**P* < 0.01.

We also observed a subpopulation of neurons that responded to N/OFQ application with a small inward current, consistent with an excitatory effect (10 nM mean response = −16 ± 6 pA) (Fig. 3A,B). Inward currents were observed in approximately 25% (15/60) of the neurons that were responsive to N/OFQ (10 nM) and 17% of all 10 nM-tested VTA neurons (15/86; 15 neurons from 14 rats; Fig. 3C,D). Among 5 neurons responding to N/OFQ with an inward current and immunocytochemically identified, 40% (2/5) were TH(+) and 60% (3/5) were TH(−) (two tailed permutation test: *p* = 0.6; Fig. 3E). These N/OFQ evoked excitatory responses were only observed at low concentrations (< 100 nM; Fig. 3D); at higher concentrations only outward currents were observed (Fig. 1E, 3D). The neurons showing this excitatory response to N/OFQ were topographically intermixed with VTA neurons that responded to N/OFQ with an outward current (Fig. 3F).

**Figure 3:**
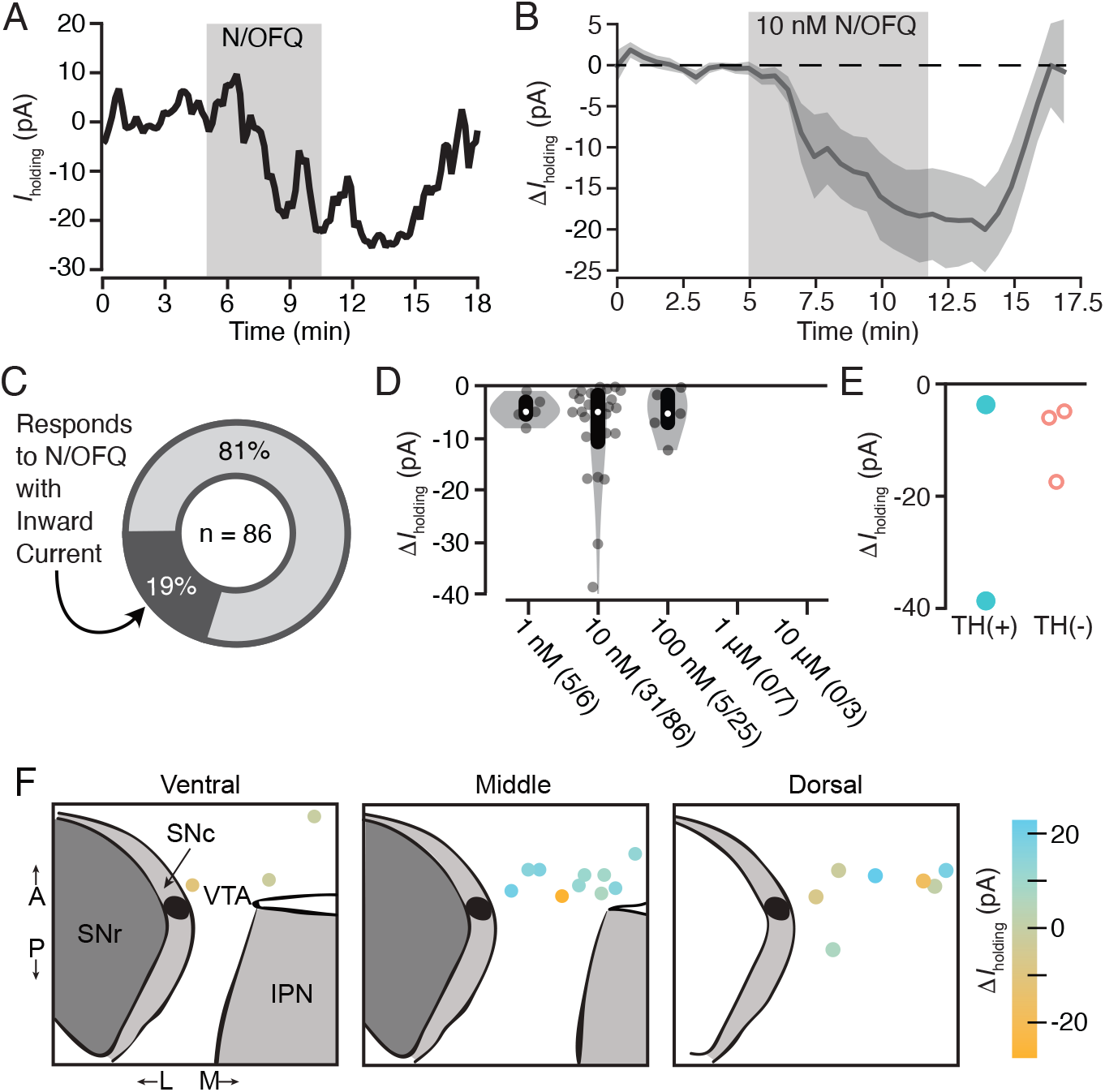
Low dose N/OFQ induced small inward currents in a subset of VTA neurons. A: Example voltage clamp recording (V_clamp_ = −60 mV) of a VTA neuron that responded to N/OFQ with an inward current. B: The mean ± SEM time course across neurons with inward currents shows the onset of this response is time locked to the initiation of drug application (n = 13). C: Across all VTA neurons from control rats that were tested for 10 nM N/OFQ responses, 19% responded with a significant inward current. D: Grey dots indicate each neuron showing a negative change, both significant and not significant, in holding current with N/OFQ application (grey dots include all neurons with a change < 0 pA; median shown in white dots; black bars show 25 and 75 percentiles. Significant inward currents were observed at 10 nM, while higher concentrations only generated outward currents (see Fig. 1E). E: Inward currents were observed in both immunocytochemically identified TH(+) and TH(−) neurons. F: Locations of VTA recordings show that neurons that responded to N/OFQ with inward and outward currents were intermixed.

### Concentration dependent desensitization of NOP

Given the inconsistencies in the reports of behavioral effects of NOP agonists and antagonists, we tested whether N/OFQ causes rapid NOP desensitization at moderate doses. We observed a concentration-dependent diminished response to a second application of N/OFQ when the first application of N/OFQ was ≥ 100 nM (n = 12 neurons from 12 rats; paired t-test *p* = 0.00003; Fig. 4A,B). This is consistent with NOP desensitization, and observed in both TH(+) and TH(−) neurons (Fig. 4B). In contrast, following administration of 10 nM N/OFQ, no difference in response was observed between the first and second applications (n = 10 neurons from 8 rats; paired t-test *p* = 0.13; Fig. 4C,D). Therefore, desensitization occurs at moderate N/OFQ concentrations in the VTA.

**Figure 4:**
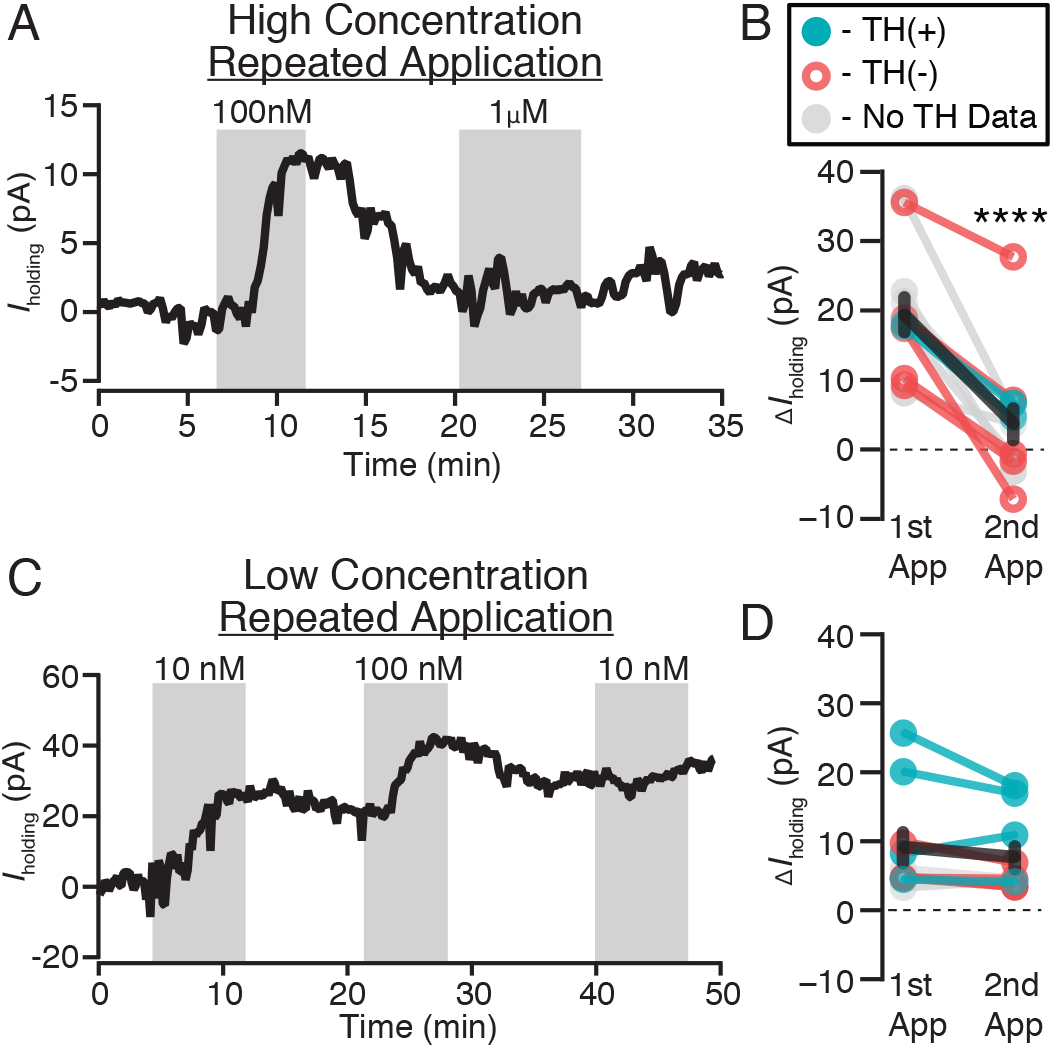
Moderate doses of N/OFQ cause functional desensitization in VTA neurons. A: Example voltage clamp recording (V_clamp_ = −60 mV) where 100 nM is sufficient to prevent a subsequent response to 1 μM application of N/OFQ. B: A summary across VTA neurons where the first N/OFQ application was ≥ 100 nM, the response to the second application was consistently smaller (\*\*\*\**p* = 0.00003), in both TH(+) and TH(−) neurons (n = 2 and 5, respectively; n = 5 no TH data). C: Example voltage clamp recording (V_clamp_ = −60 mV) showing that 10 nM N/OFQ does not impair responses to subsequent N/OFQ application. In the same cell, 100 nM did prevent additional responding. D: Summary across VTA neurons shows similar magnitudes of responses to the second application of N/OFQ when the first application was 10 nM (*p* = 0.13; TH(+) n = 4, TH(−) n = 2, TH no data n = 2).

### N/OFQ inhibits VTA neurons and SNc neurons via different cellular mechanisms

We investigated the mechanism underlying the outward currents produced by N/OFQ in VTA neurons. The most common mechanism by which Gi/o coupled receptors, including the NOP, generate somatodendritic inhibition is by activation of GIRKs. First we tested if the K^+^ channel blocker BaCl_2_ (100 μM) prevented N/OFQ induced outward currents. Surprisingly, BaCl_2_ did not prevent the outward currents induced by N/OFQ at either 100 nM (Fig. 5A) or 10 nM (Fig. 5B; one tailed permutation analysis comparing all 10 nM N/OFQ VTA observations (n = 86) to 10 nM N/OFQ observations in the presence of 100 μM BaCl_2_ (n = 7), *p* = 0.2). We next tested if a cocktail of synaptic blockers including the Na^+^ channel blocker tetrotodoxin (500 nM), the α-amino-3-hydroxy-5-methyl-4-isoxazolepropionic acid receptor (AMPAR) blocker 6,7-dinitroquinoxaline-2,3(1H,4H)-dione (DNQX; 10 μM), and the GABA_A_R antagonist bicuculline (10 μM) would alter the distribution of N/OFQ responses (Fig. 5C). Interestingly, while this cocktail did not significantly change the mean of VTA neuron N/OFQ responses (two tailed permutation analysis comparing the means of all 10 nM N/OFQ VTA observations (n = 86) to 10 nM N/OFQ observations in the synaptic blocker cocktail (n = 9), *p* = 0.16), the standard deviation of the distribution of N/OFQ responses in the presence of the inhibitor cocktail was significantly reduced, suggesting this treatment did diminish N/OFQ responses (one tailed permutation analysis comparing the standard deviations of all 10 nM N/OFQ VTA observations (n = 86) to 10 nM N/OFQ observations in the synaptic blocker cocktail (n = 9), *p* = 0.03). Since the cocktail of synaptic blockers did not yield a significant change in the mean of the responses, this indicated that both outward and inward current responses were likely diminished, and inspection of the distribution indicates that in particular the N/OFQ induced outward currents were mostly prevented by this treatment (Fig. 5C). This raised the possibilities that the outward currents are via an inhibition of AMPAR signaling, via an increase in GABA_A_R signaling, or via a non-GIRK-dependent effect of a substance released by action potential activity in the slice. We previously found that in stressed animals, DOP activation in the VTA postsynaptically increases GABA_A_R signaling in VTA neurons (Margolis et al., 2011), while spontaneous glutamate release in the VTA seems insufficient to support generating an outward current by inhibiting glutamate release (Koga and Momiyama, 2000; Margolis et al., 2005; Xiao et al., 2008). In order to test whether N/OFQ affects GABA_A_R signaling in the VTA, and whether this might account for N/OFQ induced changes in holding current, we iontophoretically applied GABA in the presence of GABABR blockade (CGP35348, 30 μM) to measure GABA_A_R responses and to bypass any potential presynaptic terminal effects. We not only found that 100 nM N/OFQ increased the amplitude of GABA_A_R responses (Fig. 5D,E), the effect on iontophoresed GABA currents was proportional to the change in holding current induced by N/OFQ (Fig. 5E), across both inward and outward currents induced by N/OFQ, making it likely that GABA_A_R signaling underlies both inward and outward currents induced by N/OFQ application to VTA neurons.

**Figure 5:**
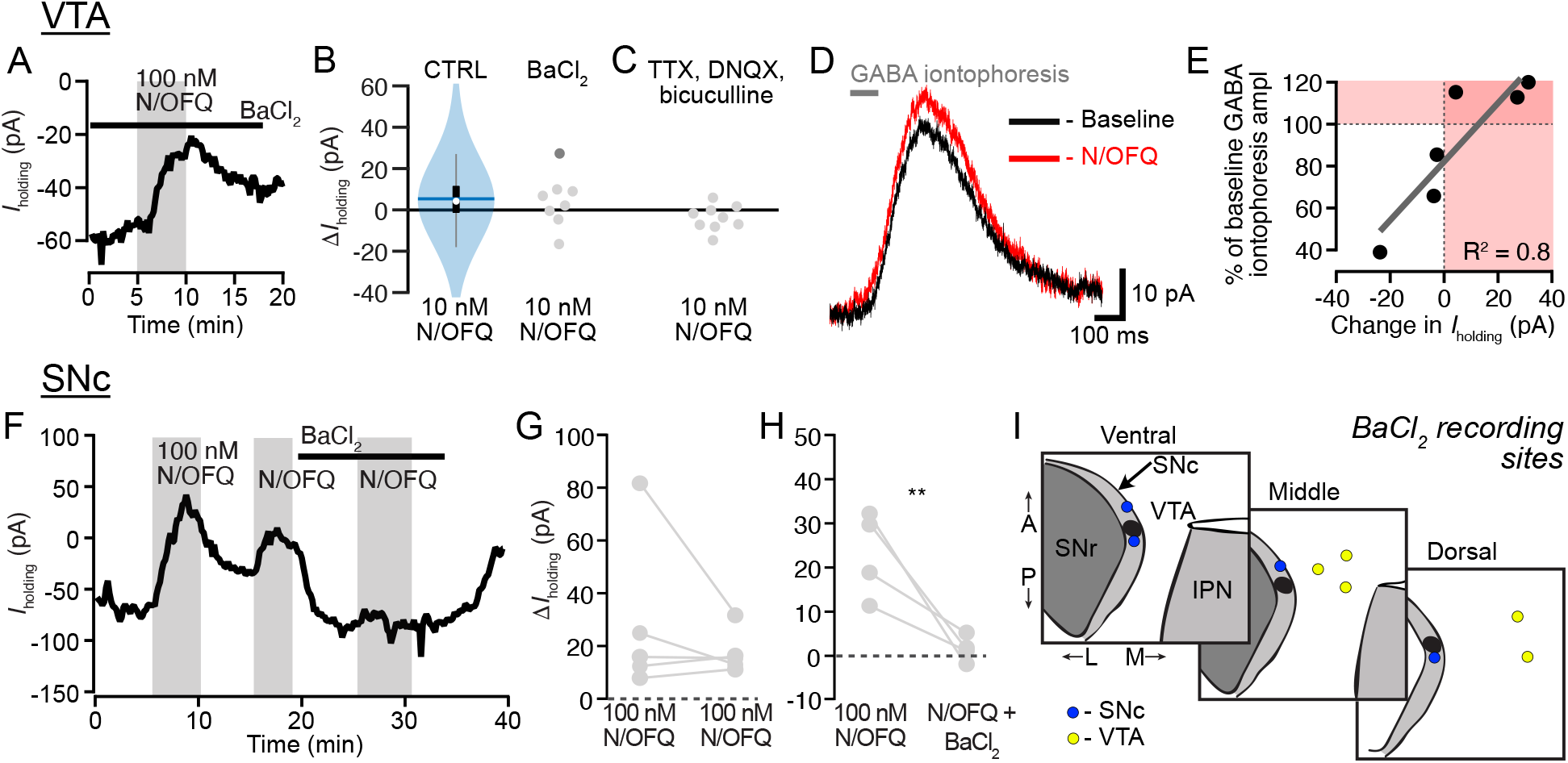
GABA_A_Rs, rather than GIRKs, mediate N/OFQ effects in VTA neurons. A: Example recording showing that the K^+^ channel blocker BaCl_2_ (100 μM) did not prevent a N/OFQ induced outward current in a VTA neuron. B: Blue violin plot represents the distribution of responses of VTA neurons to 10 nM N/OFQ (blue horizontal line = mean; white circle = median; black rectangle = 25 and 75 percentiles). In comparison, gray circles showing responses to 10 nM N/OFQ (single 100 nM experiment in dark gray) in the presence of BaCl_2_ have a similar distribution (one tailed permutation analysis of the means, *p* = 0.2). C: Recordings in 500 nM TTX, 10 μM DNQX, and 10 μM bicuculline, to block synaptic activity, AMPARs, and GABA_A_Rs, respectively, showed an almost complete elimination of outward currents in VTA neurons in response to N/OFQ (two tailed permutation analysis of the means, *p* = 0.16; one tailed permutation analysis of the standard deviations, *p* = 0.03). D: Example recording of GABA_A_R mediated iontophoretic responses to GABA (in 30 μM CGP35348 to block GABA_B_Rs), showing an augmentation of response amplitude in response to 100 nM N/OFQ. E: Summary of the N/OFQ (100 nM) induced change in iontophoretic response vs change in *I*_holding_, showing both inward and outward N/OFQ induced currents are highly correlated with N/OFQ induced changes in iontophoresis amplitude. F: Example recording in a SNc neuron showing repeated responses to high concentration (100 nM) N/OFQ, and complete blockade of the N/OFQ response by BaCl_2_. G: Summary data from SNc neurons showing minimal desensitization in control experiments with repeated within cell N/OFQ applications at high concentration. H: Summary data from SNc neurons shows that BaCl_2_ prevents a second response to N/OFQ, indicating that in the SNc, N/OFQ outward currents are mediated by K^+^ channels, unlike VTA neurons. \*\**p* < 0.01. I: Recording locations for VTA and SNc recordings where N/OFQ was tested in the presence of BaCl_2_.

That N/OFQ induced outward currents are due to augmentations of GABA_A_R mediated current rather than activation of a K^+^ current was particularly surprising because it was previously reported that N/OFQ activates a K^+^ channel in VTA neurons (Zheng et al., 2002). Zheng and colleagues also reported larger average outward currents compared to our dataset and did not observe desensitization with repeated applications of 300 nM N/OFQ, inconsistent with our findings here. As a positive control to test that 100 μM BaCl_2_ was sufficient to block K^+^ mediated effects in our preparation, and in an attempt to resolve these discrepancies, we completed additional recordings in the SNc, just lateral to the VTA (Fig. 5I). First, we tested if repeated application of 100 nM N/OFQ to SNc neurons resulted in less desensitization than we observed in VTA neurons. In fact, the response to the second 100 nM N/OFQ application was not statistically different from the response to the first application in SNc neurons, in contrast to VTA neurons (Fig. 5F,G; two-tailed paired t-Test, *p* = 0.5, n = 5). Therefore we used a within cell design to compare the N/OFQ response in control aCSF and in 100 μM BaCl_2_. Blocking K^+^ channels completely blocked the N/OFQ responses in SNc neurons (Fig. 5H; one-tailed paired t-test, *p* = 0.003, n = 5). Together, these observations indicate that BaCl_2_ was fully capable of blocking GIRK mediated N/OFQ effects in our recording conditions, and suggest that the differences between our observations and those previously reports may be related to recording location (Fig. 5I).

### N/OFQ effects on VTA neurons vary with projection target

As described above, we observed heterogeneity in responses of VTA neurons to N/OFQ. Given that other pharmacological responses of VTA neurons, including to KOP activation (Ford et al., 2006; Margolis et al., 2006) vary with projection target, we investigated whether the N/OFQ responses would be more consistent within subpopulations of VTA neurons that share a projection target. Accordingly, we recorded N/OFQ (10 nM) responses in VTA neurons that were retrogradely labeled by tracer injections into mPFC, pACC, or medial NAc (Fig. 6A,B). Recordings were conducted with the investigator blinded to the injection site.

**Figure 6:**
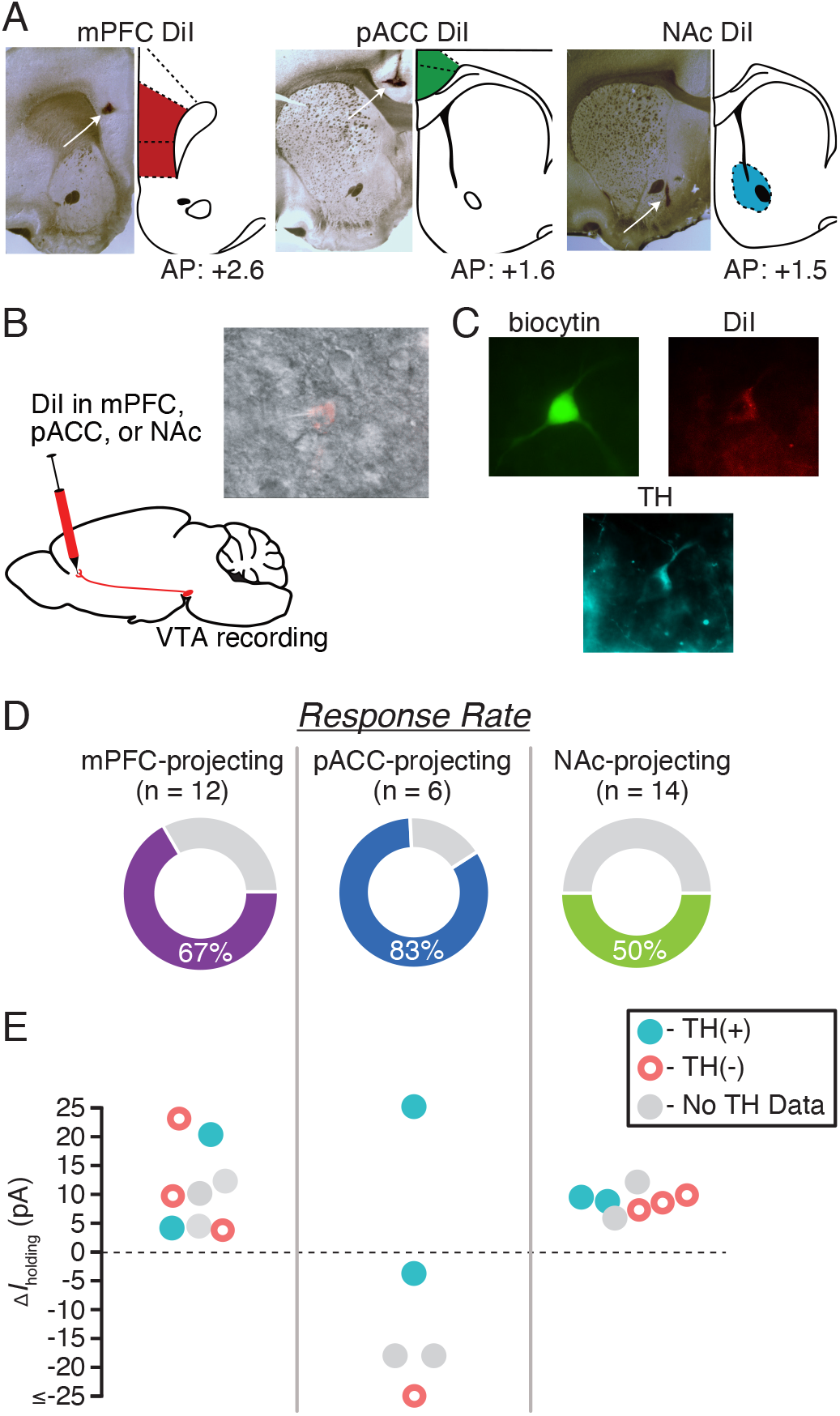
N/OFQ effects in VTA neurons vary with projection target. A: For each retrograde tracer injection site, example histology photo showing DiI localization (left) and mirrored, modified rat brain atlas schematic (right) (Paxinos and Watson, 1997). B: Cartoon showing the experimental approach: 7 days prior to recording, the retrograde tracer DiI was stereotaxically injected into mPFC, pACC, or medial NAc. DiI neurons were identified during whole cell recordings (inset). C: Example image of a neuron filled with biocytin during recording (green), the retrograde tracer (red) and was immunocytochemically identified as TH(+) (turquoise). D: The overall percentage of neurons that responded to N/OFQ was greatest among pACC-projecting neurons and lowest among NAc-projecting neurons. E: Graph of magnitudes of significant N/OFQ responses, showing that only pACC-projecting neurons respond to N/OFQ with an inward current.

The majority of mPFC-projecting VTA neurons, 67% (8/12), were significantly inhibited by N/OFQ, responding with an outward current (11 ± 3 pA; 8 responsive neurons from 6 rats; Fig. 6D). No N/OFQ induced inward currents were observed in mPFC-projecting neurons. Five mPFC-projecting neurons were recovered and processed for TH immunoreactivity (Fig. 6C,D); two were TH(+), and 3 were TH(−); all of these responded to N/OFQ with an outward current (Fig. 6D).

VTA projections to different cortical targets, including the pACC, arise from largely separate VTA neurons (Chandler et al., 2013). The pACC-projecting neurons are concentrated in different parts of the VTA, and fewer of them are dopaminergic compared to the projection to mPFC (Breton et al., 2019). Interestingly, 67% of the VTA neurons comprising this projection responded to N/OFQ with an inward current (4/6 inward current, −24 ± 12 pA, 1/6 outward current, from 4 rats; Fig. 6D). These N/OFQ excited, pACC-projecting VTA neurons included both TH(+) and TH(−) cells (Fig. 6D).

Half of NAc-projecting VTA neurons (7/14) responded to N/OFQ with outward currents (9 ± 1 pA, 7 responsive neurons from 7 rats; Fig. 6D). No inward currents were observed in this projection. Of the 7 NAc-projecting neurons that responded to N/OFQ, 2 were confirmed TH(+) and 3 were TH(−) (Fig. 6D). Together, these data indicate that similar N/OFQ inhibitory effects occur in VTA neurons that project to mPFC and NAc, but these effects are opposed to those on VTA projections to pACC, many of which responded to N/OFQ with an inward current.

### N/OFQ has little effect on terminal dopamine release in the NAc

ICV or intra-VTA N/OFQ decreases dopamine levels in the NAc (Murphy et al., 1996; Murphy and Maidment, 1999). Consistent with this result, we found that N/OFQ directly inhibits a subset of the VTA dopamine NAc-projecting somata. N/OFQ may also inhibit dopamine release in the NAc at the terminals; to test if NOPs on dopamine terminals in the NAc also contribute to an N/OFQ-induced decrease in NAc dopamine levels, we used FSCV to detect changes in stimulated dopamine release in NAc slices (Fig. 7A). In tissue from control SD rats (9 rats), we stimulated dopamine release with a bipolar electrode. In a second set of animals, to limit stimulation to dopaminergic axons, we expressed ChR2 in Th∷Cre rats and stimulated with 470 nm light pulses (9 rats). In these preparations, repeated electrical, and especially optical, stimulation can cause rundown in evoked dopamine release over time (Bass et al., 2013; O’Neill et al., 2017). To minimize this rundown as much as possible, we increased the intervals between light stimulations to 3 min. Where recordings were stable, effects of 10 nM N/OFQ, 100 nM N/OFQ, and 1 μM U69593 were sequentially tested. At 10 nM N/OFQ, approximately the EC_50_ of the outward currents recorded at VTA somata, there was no change in the peak FSCV response to either electrical or light evoked dopamine release (electrically evoked dopamine release: 93 ± 4% of baseline, n = 9 slices from 9 rats: linear mixed effects model, z = −1.3, *p* = 0.2; optically evoked dopamine release: 94 ± 10% of baseline, n = 5 from 5 rats: linear mixed effects model, z = −0.5, *p* = 0.6; Fig. 7A,B). We detected a small but significant decrease in evoked dopamine release in response to 100 nM N/OFQ (electrically evoked dopamine release: 88 ± 7% of baseline, n = 11 from 9 rats: linear mixed effects model, z = −2.1, *p* = 0.04; optically evoked dopamine release: 74 ± 4% of baseline, n = 17 from 9 rats: linear mixed effects model, z = −7.2, *p* < 0.001; Fig. 7B). Consistent with previous studies (Bass et al., 2013; O’Neill et al., 2017), it is possible that this small decrease was driven, at least in part, by rundown of ChR2-driven dopamine release. As a positive control, we applied the selective KOP agonist U69,593 (1 μM), previously shown to inhibit dopamine release in the NAc (G Di Chiara and Imperato, 1988; Ebner et al., 2010; Karkhanis et al., 2016; Spanagel et al., 1992; Werling et al., 1988), at the end of each experiment, on top of N/OFQ since these drug responses were minimal. U69,593 caused a substantial decrease in stimulated dopamine release (electrical: 53 ± 5% of baseline (in N/OFQ), n = 15: linear mixed effects model, z = −8.9, *p* < 0.001; optical: 49 ± 3% of baseline (in N/OFQ), n = 17: linear mixed effects model, z = −13.9, *p* < 0.001; Fig. 7B). Therefore, the direct NOP modulation of this dopaminergic circuit occurs at a lower concentration and may be stronger in the somadendritic region where the terminals in the NAc are relatively insensitive to NOP activation. These results contrast with the KOP control of these neurons, which strongly inhibits release at the NAc dopamine terminals but does not directly hyperpolarize the cell bodies of these neurons (Margolis et al., 2006) (Fig. 7C).

**Figure 7:**
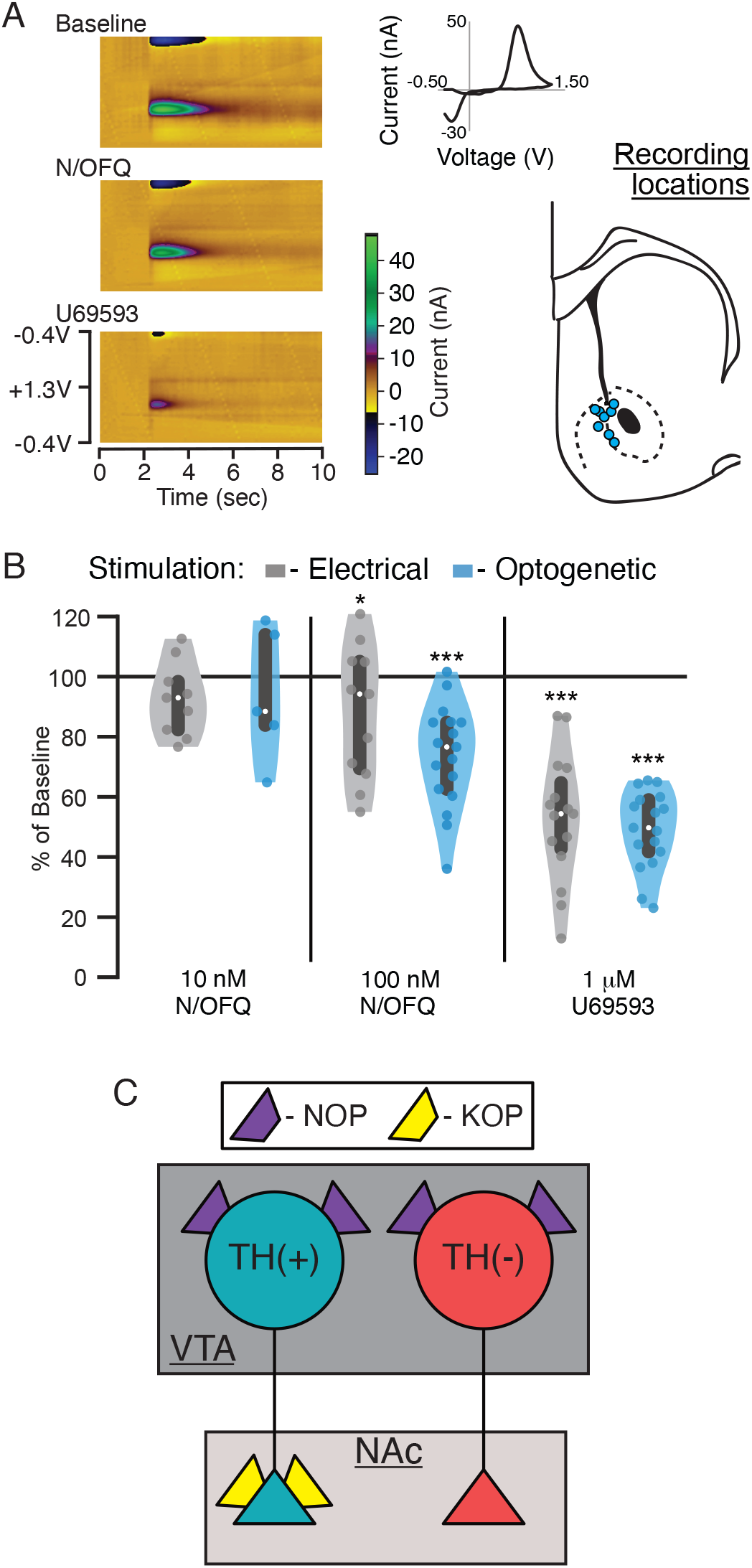
N/OFQ does not inhibit dopamine release at NAc terminals. We used FSCV in acute, coronal slices containing the NAc to test for N/OFQ effects on terminal release of dopamine. Dopamine release was evoked in slices from control rats with bipolar electrodes locally in the NAc. Recordings were made on the NAc shell-core border. Alternatively, to limit stimulation to dopamine fibers, Th∷Cre rats were injected with (AAV2-Ef1a-DIO-hChR2(H134R)-mCherry) in the VTA at least 4 weeks prior to recordings, during which 470 nm light pulses were used to stimulate dopamine release. A: Example color plots of FSCV measurement of electrically evoed dopamine release. Inset, top: background subtracted cyclic voltammogram at peak of putative dopamine release. Inset, bottom: locations of FSCV recordings in schematic of coronal section of rat brain AP: +1.5 mm (Paxinos and Watson, 1997). B: Either 10 nM or 100 nM N/OFQ was applied to the slice; each on average had minimal effects on either electrically or light evoked dopamine release in the NAc. Following N/OFQ measures, without washout, we added the KOP agonist U69593 (1 μM), which inhibited evoked dopamine release. White dots represent median values and grey bars represent 25 and 75 percentiles. C: Summary diagram shows the contrast between NOP and KOP function in NAc-projecting VTA dopamine neurons. While NOP activation inhibits the somatodendritic compartment only, KOP induced inhibition is limited to dopaminergic axon terminals in these neurons. Further, NOP activation inhibits NAc-projecting non-dopaminergic VTA cell bodies, which are insensitive to KOP activation.

## Discussion

The results presented here demonstrate that N/OFQ affects both dopaminergic and non-dopaminergic VTA neurons, through activation of the NOP, and in the majority of neurons causes inhibitory outward currents. N/OFQ effects in these neurons were blocked by the NOP selective antagonist BTRX-246040, confirming its action at NOP. Importantly, neuronal responses to N/OFQ in VTA neurons desensitized at concentrations ≥ 100 nM. In addition to providing a basic characterization of the range of postsynaptic N/OFQ responses in VTA neurons, we demonstrated differential responding of subsets of VTA neurons to NOP activation related to projection target: mPFC-projecting and NAc-projecting VTA neurons responded to N/OFQ with outward currents (inhibitory), while most pACC-projecting VTA neurons responded with inward currents (excitatory). Within the dopaminergic projection to the NAc, although N/OFQ caused outward currents at the somatodendritic region of these neurons, release at the terminals was not inhibited by NOP activation. Together, these data show that N/OFQ effects in VTA neurons differ depending upon their projection target, and that at higher concentrations of N/OFQ only inhibitions are observed, followed by desensitization of NOP function.

Unexpectedly, a small population of neurons in the VTA, both TH(+) and TH(−), responded to low concentrations of N/OFQ with an inward current, consistent with excitation. This finding presents a novel mechanism by which N/OFQ could selectively activate specific VTA circuits, while inhibiting the majority of VTA outputs. Inward currents were observed in most VTA neurons projecting to the pACC, but not those projecting to the NAc or mPFC, consistent with this circuit-selection proposition. The fact that this effect was only observed at low concentrations indicates that very robust N/OFQ release into the VTA, on the other hand, would likely have a broad inhibitory effect on the vast majority of VTA neurons, regardless of circuit. Although NOPs are generally thought to couple to Gi/o and inhibit neural activity, some exceptions to this coupling have been reported for the related opioid receptors. Activation of postsynaptic MOP or DOP results in a Ca_v_2.1 channel-dependent depolarization in subsets of VTA neurons (Margolis et al., 2017, 2014). Further, the MOP agonist DAMGO increases Ca_v_2.1 currents in cerebellar Purkinje neurons (Iegorova et al., 2010) and morphine activates adenylyl cyclase in the corpus striatum and olfactory bulb (Onali and Olianas, 1991; Puri et al., 1975). While this is the first report of N/OFQ-mediated excitations in an acute brain slice preparation, intracellular increases in Ca^2+^ have been observed in a cultured human neuroblastoma cell line in response to N/OFQ in the presence of the cholinergic agonist carbachol (Connor et al., 1996). Therefore, while there are few reports of excitatory actions of N/OFQ, the observation is not unprecedented.

We also found that NOP activation signals through the canonical GIRK pathway in the SNc, however, in the VTA N/OFQ outward currents were mediated by augmentation of GABA_A_R currents. While both mechanisms generated outward currents in our experimental preparation, the physiological consequences of these neural populations utilizing different signaling pathways *in vivo* may vary. For instance, activating a GIRK will always cause a hyperpolarization, while increasing the GABA_A_R current will only occur when there is concurrent activation of NOPs and GABA_A_Rs. Further, the N/OFQ induced inhibition requiring GABA_A_R activation depends upon the Cl^−^ reversal potential, which may be altered by a variety of behavioral states including pain, morphine treatment, stress, or alcohol (Coull et al., 2003; Ferrini et al., 2013; Hewitt et al., 2009; Ostroumov et al., 2016; Santos et al., 2017). The N/OFQ response may even be excitatory in the absence of GABA_A_R activation, since blocking GABA_A_Rs seemed to increase the proportion of neurons in which we observed inward currents in response to N/OFQ (Fig. 5C).

In the VTA, neurons treated with a higher concentration of N/OFQ (≥ 100 nM) no longer responded to subsequent applications of N/OFQ in the VTA. This finding indicates that N/OFQ may act as a functional antagonist at the NOP by desensitizing these responses when higher concentrations of N/OFQ are present. Interestingly, we did not observe significant NOP desensitization in SNc neurons. NOP function is therefore apparently different from postsynaptic responses to agonists at the MOP and DOP in VTA neurons, where repeated application of saturating concentrations of selective agonists generate responses of similar magnitudes (Margolis et al., 2017, 2014). The apparent NOP desensitization we observed in the VTA is consistent with previous studies showing that high concentrations or repeated sustained exposure to NOP agonists causes desensitization in cell culture (Connor et al., 1996; Mandyam et al., 2002, 2000; Thakker and Standifer, 2002). In addition, NOPs internalize fairly rapidly (Corbani et al., 2004; Spampinato et al., 2002, 2001; Zhang et al., 2012) at the same concentrations that we observed desensitization. *In vivo*, N/OFQ administration can result in dose dependent performance changes in behavioral spatial memory, locomotor, and anxiety tasks, with low concentration N/OFQ having the opposite effects compared to high doses (Florin et al., 1996; Jenck et al., 1997; Sandin et al., 2004). One possible explanation for these opposing behavioral outcomes is that N/OFQ may be acting as an agonist at low concentrations and a functional antagonist at high concentrations in some brain regions. An alternative possibility is that brain regions like the SNc that have less desensization drive behavioral responses to high doses of N/OFQ, where brain regions like the VTA that show more desensitization do contribute to behavioral responses to lower N/OFQ doses.

N/OFQ inhibited both dopamine and non-dopamine neurons in the VTA that project to the NAc. This finding is consistent with the observation that N/OFQ administered ICV or into the VTA results in a decrease in extracellular dopamine in the NAc (Murphy et al., 1996; Murphy and Maidment, 1999). A prominent proposal in the literature is that a decrease in NAc dopamine produces aversion (McCutcheon et al., 2012). Therefore, one would expect ICV injection of N/OFQ to be aversive. However, this manipulation generates no response in the place conditioning paradigm (Devine et al., 1996). On the other hand, optogenetic or chemogenetic stimulation of N/OFQ containing inputs to the VTA can be aversive and decrease reward seeking (Parker et al., 2019). One possible explanation for this lack of clear motivational effect is the combination of inhibition of both dopamine and non-dopamine neurons: dopamine and non-dopamine neurons originating in the VTA synapse onto different types of neurons in the NAc, therefore affecting behavior in different ways. For instance, VTA glutamate neurons synapse onto parvalbumin containing interneurons in the NAc and optogenetic activation of these NAc-projecting glutamate neurons is aversive (Qi et al., 2016). Activation of NAc-projecting VTA GABA neurons causes a pause in cholinergic interneuron activity (Brown et al., 2012). These neurons modulate associative learning but are insufficient to drive preference or aversion independently (Collins et al., 2019) and do not appear to contribute to the detection of aversive gustatory stimuli (Robble et al., 2020). N/OFQ inhibition of dopamine, GABA, and glutamate neurons projecting to the NAc, therefore, may result in no net hedonic value and a lack of preference in a place preference paradigm. Further, various reports indicate that decreasing activity at dopamine receptors in the NAc with microinjections of antagonists does not produce aversion (Baker et al., 1998, 1996; Fenu et al., 2006; Josselyn and Beninger, 1993; Laviolette and van der Kooy, 2003; Morutto and Phillips, 1998; Spina et al., 2006) but see (Shippenberg et al., 1991), and aversive outcomes can even be observed following manipulations that increase dopamine levels in the NAc (Devine et al., 1993b; Shippenberg and Bals-Kubik, 1995). Add to this the N/OFQ effects on other circuits following ICV injection, including other VTA neurons, and the possibility that the most robust, long lasting effect is receptor desensitization at higher doses of agonist, together making it potentially less surprising that ICV N/OFQ was not reported to generate aversion.

N/OFQ’s effect on the VTA to NAc circuit provides an interesting point of comparison for how the NOP may be functionally distinct from the structurally related KOP. *In vivo*, systemic or ICV administration of N/OFQ or a KOP agonist each causes a decrease in extracellular dopamine in the NAc (Devine et al., 1993a; G. Di Chiara and Imperato, 1988; Murphy et al., 1996; Murphy and Maidment, 1999). However, these two receptors function very differently in the dopamine neurons that project to the NAc. We show here that N/OFQ inhibits VTA cell bodies that project to the NAc, but has little effect on the dopamine terminals within the NAc. Kappa opioid receptor activation, on the other hand, has no effect on the cell bodies of NAc-projecting VTA dopamine neurons, but strongly inhibits dopamine release at the terminals in the NAc (Britt and McGehee, 2008; Margolis et al., 2006) (Fig. 5C). One implication for this organization is that whether or not the respective endogenous peptides, N/OFQ and dynorphin, affect NAc-projecting dopamine neurons will depend upon the brain region of peptide release. There is also evidence for dopamine release in the NAc that is independent of action potential firing in midbrain dopamine neurons (Cachope et al., 2012; Mohebi et al., 2019). In this organization of differential receptor effects localized to somadendritic regions vs terminals, dynorphin has control over this terminal activity while N/OFQ does not. Together these observations bring into focus the critical importance of understanding precisely where receptors are functional in brain circuits and their specific actions at each site.

We found opposing effects of N/OFQ on the VTA projections to mPFC and pACC, which may contribute to the reported N/OFQ impact on behavioral measures associated with cortical dopamine function such as working memory, learning, and behavioral flexibility (Gonzalez et al., 2014; Huang et al., 2018; Ott and Nieder, 2019; Puig et al., 2014; Tzschentke, 2001; Winter et al., 2009). Our results also show that the non-dopamine VTA projection to cortical regions are affected by N/OFQ as well; while the majority of the VTA neurons that project to these cortical regions are not dopaminergic (Breton et al., 2019), little is currently known regarding their contribution to behavior. Preclinical studies show that ICV administration of N/OFQ impairs working memory (Hiramatsu and Inoue, 1999) and associative learning and memory (Goeldner et al., 2009), and blocking NOP with an antagonist or genetic knockout enhances both working memory and learning (Jinsmaa et al., 2000; Nagai et al., 2007; Noda et al., 1999). How and why such an ongoing break on learning and memory by N/OFQ contributes to normal behavioral adaptation, and whether dopamine or other VTA outputs play a role, remains to be determined. One provocative possibility is that it is this degradation of working memory function that is the primary mechanism underlying the lack of place conditioning in response to central N/OFQ, rather than that this treatment is affectively neutral. This interpretation is consistent with work showing that N/OFQ blocks opioid induced conditioned place preference yet has no effect on opioid self-administration as well (Sakoori and Murphy, 2004; Walker et al., 1998).

The results of this study extend our understanding of the NOP system biology and provide considerations for additional investigation into NOP function within limbic circuits. These findings clarify that strong NOP desensitization occurs in neurons at moderate concentrations of the endogenous agonist N/OFQ. Importantly, not only does the nature of the NOP response vary with the projection target of VTA neurons, but the NOP function is largely sequestered to the somatodendritic compartment of VTA dopamine neurons that project to the NAc, demonstrating two different kinds of circuit level organization of this receptor system. Building on this groundwork, future studies of these VTA circuits during different behavioral states and tasks related to motivation and cognition will help to elucidate the differences between the normal and dysfunctional NOP-N/OFQ system, improving the potential for therapeutic targeting.

---

**Extended Data:**
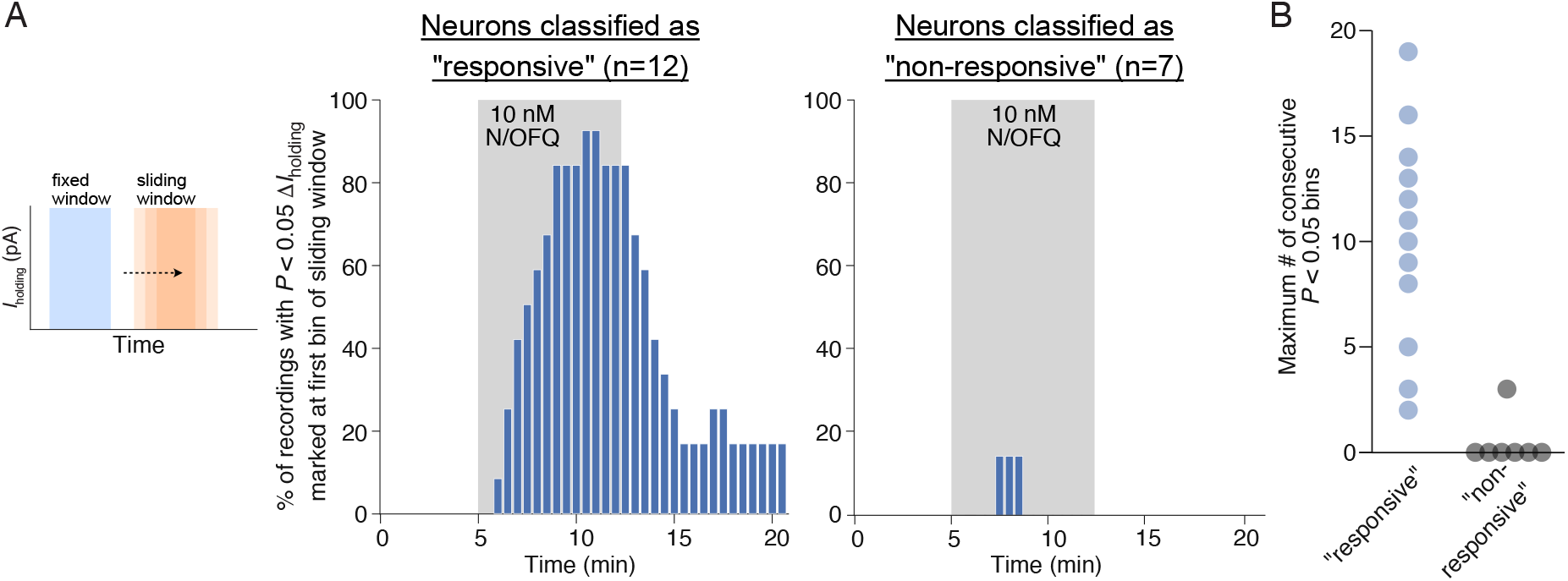
To evaluate our within cell statistical comparisons to identify “responsive” vs “non-responsive” neurons, in particular to test the possibility that drift might contribute to some of our identified drug effects, we conducted a sliding window analysis on a subset of our drug responses (all TH positive neurons tested with bath application of 10 nM N/OFQ). Further, any increase in statistically significant sliding windows during drug washout compared to the static baseline would suggest underlying *I*_holding_ drift. We compared all 4 minute windows from pre-drug application through drug washout to a fixed “baseline” window (the 4 min preceding the onset of the drug). To create the windows, *I*_holding_ of each recording was binned into 30 second intervals and assigned a bin number (1, 2, 3 Ö n). The “baseline” 8 bin (4 min) window was compared with the target 4 min window by way of a student’s unpaired t-test. The *P* value and significance of the comparison was then corrected using the Bonferroni method for multiple comparisons. The alignment of the sliding window was then increased by a single bin and the comparison repeated, resulting in an array that represents all significant 4 minute intervals for each drug effect. The resulting arrays were plotted as a histogram representing, at the initial bin time of the sliding window, the proportion of recordings in which this calculation was significantly different from the fixed baseline target window (A). In the neurons previously classified as “responsive” by a single “baseline” compared to “drug” window comparison, the rising left edge of the histogram begins to plateau around the 4^th^ minute of drug application, consistent with the plateau of the mean effects across all cells reported in Figure 1D. Further, consistent with washout reversal of N/OFQ effects in most but not all neurons, the proportion of significant bins falls off as soon as N/OFQ application was terminated. That both the rise and fall of the frequencies of significant windows are time locked to the drug application suggests the response classification scheme is reliable. In neurons previously classified as “non-responsive” only one neuron had any significant windows, with 3 sliding window locations where this analysis yielded *P* < 0.05, suggesting that there was not systematic drift in these “non-responsive” neurons. In addition, a scatter plot (B) indicates the maximum number of consecutive significant sliding windows for each cell analyzed, because a well-behaved change in *I*_holding_ in response to the drug application should be detected in consecutive sliding windows. This graph shows that 8/12 neurons that were classified as “responsive” have more consecutive sliding windows different from baseline than the maximum found in “non-responsive” neurons. This analysis was conducted using a custom script created in Python (available at https://osf.io/c8gu7/?view_only=63ea4c0623b54e46a4efaccc450a89c6).

## Acknowledgements

We would like to thank Benjamin J. Snyder and Venkateswaran Ganesh for their technical assistance. We also thank Howard L. Fields for helpful comments on the manuscript.

